# Leafminer attack induces plant-mediated facilitation of conspecific pupae in the soil

**DOI:** 10.1101/2020.10.07.329763

**Authors:** Rocío Escobar-Bravo, Bernardus CJ Schimmel, Gaétan Glauser, Peter GL Klinkhamer, Matthias Erb

## Abstract

- Herbivore population dynamics are strongly influenced by the interactions established through their shared host. Such plant-mediated interactions can occur between different herbivore species and between different life developmental stages of the same herbivore. Yet, whether these interactions occur between leaf-feeding herbivores and their soil-dwelling pupae is unknown.
- We studied whether tomato (*Solanum lycopersicum*) leaf-herbivory by the American serpentine leafminer *Lyriomiza trifolii* affects the performance of conspecific pupae in the soil adjacent to the plant. To gain mechanistic insights, we performed insect bioassays with the jasmonate-deficient tomato mutant *def-1* and its wild type, along with the analysis of phytohormones, gene expression and root volatiles.
- Leafminer metamorphosis in the soil was accelerated when wild type plants were attacked aboveground by conspecifics, but the opposite was observed in *def-1*. Changes in pupal developmental rate were mediated by belowground volatiles. Accordingly, leafminer herbivory differentially modulated jasmonate and abscisic acid signaling and the accumulation of specific volatiles in the roots of wild type versus *def-1* plants
- Our results demonstrate that aboveground herbivores can facilitate their soil-dwelling pupae by inducing *def-1*-dependent systemic responses. This study expands the repertoire of plant-herbivore interactions to herbivory-induced modulation of metamorphosis, with potentially important consequences for plant and herbivore community dynamics.

## Introduction

Plants are attacked by multitudes of herbivorous arthropods (Stam *et al.*, 2014). Upon attack, plants can alter their primary and secondary metabolism to produce toxic or anti-nutritive substances that reduce herbivore preference, performance and/or survival (Mithöfer & Boland, 2012). These chemical readjustments are controlled by plant hormonal signaling, among which the jasmonate pathway is a central regulator (Erb & Reymond, 2019). Some herbivores, however, have developed intricate strategies to suppress or modify induced plant responses to their own advantage (Sarmento *et al.*, 2011; Robert *et al.*, 2012b; Chung *et al.*, 2013). Herbivores population dynamics are therefore strongly influenced by the interplay between the induction of plant chemical defenses and herbivores manipulation of these defenses (van Dam & Heil, 2011; Soler *et al.*, 2013; Kant *et al.*, 2015).

Plant defense induction and manipulation can influence the interactions between the plant and different life stages of the herbivore (Erwin *et al.*, 2014). For instance, many herbivorous species complete their life cycle in different organs of the host plant, e.g., adults feeding on shoots and conspecific larvae feeding on roots. Initial leaf herbivory by adults can elicit systemic plant responses in the roots that facilitate or inhibit subsequent larval performance (Erwin *et al.*, 2014; Huang *et al.*, 2014; Kraus & Stout, 2019). While the impact of herbivory-induced plant defense responses on different plant-feeding life stages of conspecific and heterospecific herbivores have been studied extensively, little is known about their influence on non-feeding life stages such as the pupal stage. This is surprising considering the large number of herbivorous arthropod species (including insects belonging to the orders Diptera, Hymenoptera, Thysanoptera, Coleoptera and Lepidoptera) that pupate in the soil close to their host and, thereby, their potential exposure to the chemicals released by the roots and their associated rhizosphere. In fact, as insect pupae are immobile, their survival, i.e. adult emergence, is highly influenced by the soil habitat (Torres-Muros *et al.*, 2017).

As drivers of soil biotic and abiotic properties and producers of biologically active metabolites, plant roots play a key role in plant-mediated interactions between leaf- and root-feeding herbivores and their life stages (van Dam, 2009; Robert *et al.*, 2012a). Leaf herbivory can modulate the release of organic chemicals from roots, thus shaping the surrounding habitat and the plant-associated soil communities (Barber *et al.*, 2014; Huang *et al.*, 2017, Karssemeijer *et al.*, 2020). In particular, root emission of volatile organic compounds (VOCs) can influence the behavior of root herbivorous insects and their natural enemies (Rasmann *et al.*, 2005; Robert *et al.*, 2012b; Huang *et al.*, 2017) and potentially affect the physiology of the herbivore (Veyrat *et al.*, 2016; Ye *et al.*, 2018). Insect pupae can exchange gases through their spiracles (Jõgar *et al.*, 2007; Nestel *et al.*, 2007), it is thus conceivable that leaf attack by herbivores may influence pupal development in the soil through systemic changes in the release of root volatile chemicals.

The American serpentine leafminer *Lyriomiza trifolii* (Burgess) (Diptera, Agromyzidae) is a polyphagous insect herbivore that causes large economic losses of ornamental plants and vegetable crops worldwide (Minkenberg *et al.*, 1986). Adults of *L. trifolii* are small flies (ca. 2 mm long). Females’ flies feed and lay eggs inside the leaf, below the epidermis. The larvae subsequently feed within the leaf creating tunnels (mines). At the end of their third developmental stage, the larvae leave the mine and drop to the soil to pupate (Parrella *et al.*, 1983). Due to *L. trifolii*’s rapid life cycle (12-24 days), plants infested with this leafminer can generally hold several generations of this insect, which means that adults, larvae, and soil-dwelling pupae simultaneously occur in the field. *L. trifolii* is an important pest of cultivated tomatoes (*Solanum lycopersicum*) and can build up to high population densities during the growing season (Abe & Kawahara, 2001). Tomato roots are reported to produce and emit VOCs that influence the behavior of soil-living nematodes (Murungi *et al.*, 2018). The *L. trifolii*-tomato system is thus an ideal model to investigate if and how plants mediate interactions between leaf-feeding herbivores and their soil-dwelling pupae.

Very little is known about the interactions between *L. trifolii* and cultivated tomato at the molecular level (Stout *et al.*, 1994). Previous work in Arabidopsis suggests that plant resistance to *L. trifolii* mainly depends on jasmonate-regulated defense responses (Abe *et al.*, 2013). Here we performed insect bioassays to investigate whether *L. trifolii* aboveground infestation of tomato plants affect the performance of the leafminer pupae in the soil through changes in belowground volatiles. In addition, we investigated if these interactions depend on intact defense signaling through the use of the tomato mutant *defenseless-1* (*def-1*), deficient in the biosynthesis of jasmonic acid in response to wounding or herbivore attack (Li *et al.*, 2002). To gain insight into the chemical and molecular mechanisms underlying the tomato-leafminer interaction, we surveyed the accumulation of defense-related phytohormones, gene transcripts and root volatiles. Our results show that *L. trifolii* pupae develop faster when exposed to belowground volatiles from leafminer infested tomato plants, and that these interactions are *def-1* dependent and partially explained by the induction of JA associated responses. Furthermore, *L. trifolii* infestation significantly altered the production of tomato root volatiles in a genotype-dependent manner. Together, our results indicate that leafminer facilitation of conspecific pupae in the soil is plant-mediated, and that JA and root volatiles play a relevant role in this interaction. These findings uncover herbivory-induced modulation of metamorphosis as a new phenomenon in plant-insect interactions.

## Materials and Methods

### Plant and insect materials

Tomato (*Solanum lycopersicum* Mill.) seeds of the jasmonate-deficient mutant *defenless-1, def-1,* and its wild type, cv ‘Castlemart’, were sown in plastic trays filled with potting soil in a climate room provided with 113.6 μmol·m^2^os^-1^ of photosynthetically active radiation (PAR), a photoperiod of 16L:8D, 20°C, and 70% RH. Fifteen days after germination plantlets were transplanted to 13×13-cm plastic pots filled with the same potting soil.

A colony of the American serpentine leafminer *Lyriomiza trifolii* (Burgess) (Diptera: Agromyzidae) was maintained on chrysanthemum plants in a climate room at 25°C and 70% RH.

### Experimental design

Five-week-old wild type and *def-1* plants were individually placed inside transparent plastic cylinders (80 cm height, 20 cm diameter) closed at one end with a lid made of insect-proof gauze (Leiss et al. 2009) (Fig. 1a). Seven *L. trifolii* adult female flies were released inside each cage. Non-infested enclosed plants served as controls. Aboveground *L. trifolii* infested- and non-infested plants were randomly placed in a climate room provided with 113.6 μmol·m^-2·^s^-1^ of PAR, a photoperiod of 16L: 8D, 25°C and 70% RH. Aboveground infested plants were damaged by feeding and ovipositing *L. trifolii* adult flies and, subsequently, by the developing larvae that create tunnels (mines) within the leaves. Eight days after infestation a custom-made cage containing fifteen age-synchronized *L. trifolii* pupae (obtained from the rearing maintained on chrysanthemum; see above) was buried into the soil (~2 cm deep) close to the stem of each plant (Fig. 1a). Custom-made cages consisted of a petri-dish, sealed with parafilm, whose bottom lid was perforated and covered with an air-permeable but insect-proof mesh (Fig. 1a). Before the start of aboveground *L. trifolii* infestation treatment, above- and below-ground compartments of the plant were separated by covering the soil with aluminum foil. This system therefore exposed the pupae to belowground VOCs only. The number of emerged adults per cage and plant was then recorded every two days for a period of 16 days. At the end of the experiment, i.e., 24 days after infestation, the number of *L. trifolii* larvae-associated mines per plant and soil humidity were determined. In a second repetition of the experiment, the effects of aboveground *L. trifolii* infestation on plant hormone concentrations, gene expression levels, root volatile contents, and root dry biomass were determined at 24 days after infestation, i.e., 16 days after pupation (see methods below). Note that experiments with wild type and *def-1* plants were carried out simultaneously, and the data were statistically analyzed as such.

**Figure 1.**
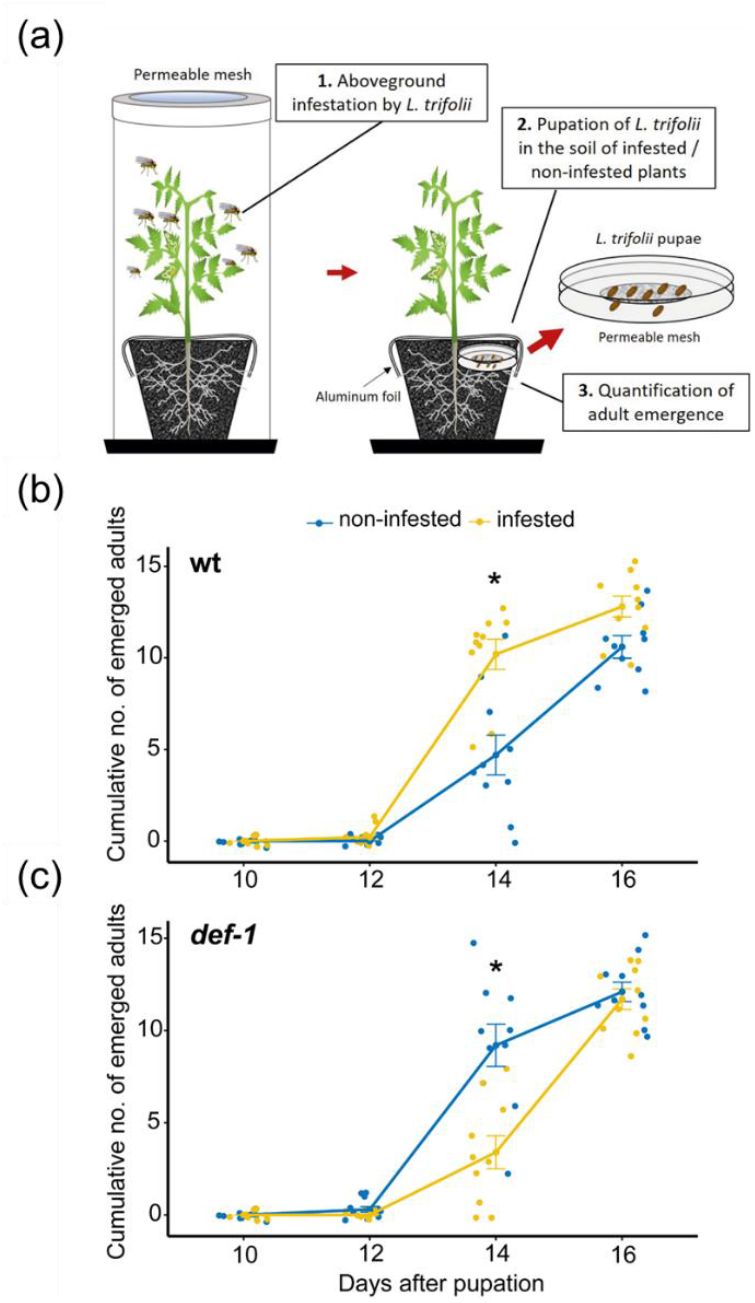
Leafminer attack facilitates the development of conspecific pupae in the soil in a *def-1* dependent manner. (a) Overview of the set-up to test the effect of *L. trifolii* leaf herbivory on the performance of conspecific pupae in the soil. Five-week-old tomato plants were individually placed inside transparent plastic cylinders closed at one end with a lid made of insect-proof gauze and infested with seven *L. trifolii* adult female flies allowing them to feed and deposit eggs on the plants. Prior to *L. trifolii* infestation, the soil was covered with aluminum foil to separate the belowground and aboveground plant compartments. Eight days after infestation, fifteen age-synchronized *L. trifolii* pupae from an external rearing were confined to a custom-made and air-permeable cage and subsequently buried into the soil of each plant. Cumulative number of *L. trifolii* adults (mean ± SEM, *n* = 10) that emerged from pupae exposed to belowground volatiles from (b) non-infested or *L. trifolii*-infested wild type (wt) plants and (c) non-infested or *L. trifolii*-infested *def-1* plants were recorded every two days for a total of 16 days. Data from panels (b) and (c) were obtained from the same experiment and statistically analyzed altogether by using a negative binomial model that included plant genotype, herbivory and their interactions as fixed effects, and plant unit as the random intercept. Asterisks denote significant differences tested by pairwise comparisons of estimated marginal means with the Tukey-adjusted *P* value method (*p* ≤ 0.05).

### Soil humidity

Soil humidity was estimated using the gravimetric method (Parkin *et al.*, 2008). A sample of fresh soil per plant was placed in a 2 ml Eppendorf tube, weighed, oven dried at 60°C for 5 days, and re-weighed. Fresh and dry soil mass (g) were calculated by subtracting the weight of empty 2 ml Eppendorf tubes. Gravimetric water content (GWC) expressed as the mass of water per mass of dry soil was then calculated as follows: GWC = [*wet soil* (*g*) – *weight dry soil* (*g*)] / *weight of dry soil* (*g*).

### Phytohormone analysis

The concentrations of 12-oxo-phytodienoic acid (OPDA), JA, jasmonic acid-isoleucine (JA-Ile), salicylic acid (SA), and abscisic acid (ABA) were determined in the third leaf from the apex and the roots of non-infested and infested wild type and *def-1* plants at 24 d after aboveground *L. trifolii* infestation. For this, the phytohormones were extracted with ethyl acetate spiked with isotopically labeled standards (1 ng of d5-JA and 13C6-JA-Ile, d4-SA, and d6-ABA) and analyzed by ultra-high-performance liquid chromatography tandem mass spectrometry (UHPLC-MS/MS) following the procedures described in Glauser et al. (2014).

### Gene expression analysis

Total RNA was isolated using TRI Reagent (Sigma-Aldrich) following the manufacturer’s instructions. DNase treatment was carried out using gDNA Eraser (Perfect Real Time) following manufacturer’s instructions (Takara Bio Inc. Kusatsu, Japan). Reverse transcription and first strand cDNA synthesis was performed using PrimeScript Reverse Transcriptase (Takara Bio Inc.). For gene expression analysis, 2 μl of cDNA solution (the equivalent to 10 ng of total RNA) served as template in a 10 μl qRT-PCR reaction using ORA™ SEE qPCR Mix (Axonlab) on an Applied Biosystems® QuantStudio® 5 Real-Time PCR System. The normalized expression (NE) values were calculated with the ΔCt method: NE = (1/(PE_target_^Ct_target_))/(1/(PE_reference_^Ct_reference_)) in which PE stands for primer efficiency and Ct for cycle threshold, as in Alba *et al.* (2015). The PEs were determined by fitting a linear regression on the Ct-values of a standard cDNA dilution series. Gene identifiers, primer sequences and references are listed in Table S1.

### Root volatiles analysis

Root volatiles from *L. trifolii* infested and non-infested wild type and *def-1* plants were collected and analyzed using solid-phase micro-extraction-gas chromatography-mass spectrometry (SPME-GC-MS). For this, the whole root system was sampled, washed with tap water, gently dried with filter paper, and flash-frozen in liquid nitrogen prior to analysis. Roots were then ground into a fine powder, and 100 mg was placed in a 20 ml precision thread headspace glass vial sealed with a septum of silicone/ PTFE and a screw cap made of magnetic silver with a hole (Gerstel GmbH & Co. KG). An SPME fiber (100 μm polydimethylsiloxane coating; Supelco, Bellefonte, PA, USA) was then inserted through the screw cap hole and septum for volatile collection for 40 min at 50°C. Collected volatiles were thermally desorbed for 3 min at 220°C and analyzed by GC-MS (Agilent 7820A GC interfaced with and Agilent 5977E MSD, Palo Alto, CA, USA). Desorbed volatiles were separated with split-less mode on a column (HP5-MS, 30 m, 250 μm ID, 2.5-μm film, Agilent Technologies, Palo Alto, CA, USA) with He as carrier gas and at a flow rate of 1 mL min^-1^. A temperature gradient of 5°C min ^-1^ from 60°C (hold for 1 min) to 250°C was used. Compound identification was based on similarity to library matches (NIST search 2.2 Mass Spectral Library. Gaithersburg, MD, USA) as well as retention time and spectral comparison with pure compounds.

### Statistical analysis

Data analysis was conducted in R version 4.0 (R core Team, 2016). We modeled the cumulative number of emerged adults with a negative binomial model using *glmmTMB* package (Brooks *et al.*, 2017). Day of emergence, i.e., 12, 14 and 16 days (days where there were zero emerged adults for all treatments were not included in the analysis), plant genotype, herbivory and their interactions were included in the model as fixed effects, and plant unit as the random intercept. Residuals diagnosis, zero inflation and over dispersion tests were performed using the *DHARMa* package (Hartig, 2017). The significance of the fixed effects from the conditional model was tested via ANOVA-type analysis (type II Wald Chi-square tests). Estimated marginal means (EMMeans), standard errors, confidence limits and significant differences between means with multiple comparison adjustments (Tukey’s HSD) were estimated. Effects of herbivory, plant genotype and their interaction on hormone levels, gene expression, soil humidity, root dry biomass, and levels of root volatile compounds were tested by two-way ANOVAs. Pairwise comparisons of EMMeans with multiple comparison adjustments (Tukey’s HSD) was used as post hoc test. Prior to analysis, all residuals were tested for normality and equal variance using the Shapiro-Wilk and Levene tests, respectively. Data that did not meet these assumptions were log10, squared root, or Tukey’s Ladder of Powers transformed prior to statistical analysis. The correlation between the cumulative number of emerged adults at 14 or 16 days after pupation and (1) the number of larvae per plant or (2) the soil humidity was determined by Pearson product moment and Spearman’s rank correlation tests. Principal component analysis (PCA) was conducted to explore root volatile profile differences among treatments. For this, data were first log (x+10) to correct for the presence of zeros and skewness, the means were centered to equals to zero, and the variables were scaled to have standard deviation equals to 1. To identify which factors significantly influenced the observed chemical variation in PCA we subsequently performed a redundancy analysis (RDA) followed by permutation *F*-test (999 permutations) based on the canonical *R^2^* (Hervé *et al.*, 2018). Finally, we conducted permutation *F-*test (999 permutations) to assess the influence of plant genotype, herbivory, and their interaction on the chemical data, as well as the differences among groups by pairwise comparisons corrected with the false discovery rate method. These statistical analyses were performed and/or visualized using the R packages “*car*”, “*olsrr*”, “*rcompanion*”, “*nlme*”, “*emmeans*”, “*ggplot2*”, “*ggpbur*”, “*factoextra*”, *“Hotelling”, “vegan”,* and *“RVAideMemoire”* (Wickham & Wickham 2007; Oksanen *et al.*, 2013; Hervé, 2015; Fox *et al.*, 2016; Hebbali & Hebbali, 2017; Kassambara & Mundt, 2017; Mangiafico & Mangiafico, 2017; Pinheiro *et al*., 2017; Curran, 2018; Lenth, 2019; Kasambara, 2020)

## Results

### Leafminer herbivory facilitates pupal development in the soil

To test if aboveground infestation by the leafminer *L. trifolii* affects the performance of conspecific pupae in the soil, we infested wild type tomato plants with *L. trifolii* and monitored pupal development over time in the soil of infested and non-infested plants (Fig. 1a). To monitor adult emergence, age-synchronized *L. trifolii* pupae were placed in Petri dishes and subsequently buried in the soil 8 days after aboveground leafminer infestation. The Petri dishes were fitted at the bottom lid with a permeable nylon mesh that allowed for exchange of volatile chemicals between the rhizosphere and the pupae. Fourteen days after pupation, two times more adults emerged from batches of pupae that had been exposed to belowground volatiles from infested plants compared to non-infested plants (Fig. 1b, Table S3 and Fig. S1, day 14, wild type non-infested vs infested *p* ≤ 0.001, EMMeans pairwise comparisons). This difference was not significant at the end of the emergence period (Fig. 1b, Table S3 and Fig. S1, day 16, wild type non-infested vs infested *p* > 0.05, EMMeans pairwise comparisons). Thus, aboveground *L. trifolii* infestation accelerates the pupation of conspecifics in the soil, an effect that is most likely mediated by belowground volatile chemicals.

### Facilitation of pupal development is *def-1* dependent

Induced plant defenses, including those regulated by jasmonates, can mediate systemic shoot-root responses upon herbivore attack (Machado *et al.*, 2018). To gain insights into the potential involvement of JA-mediated signaling in leafminer-induced facilitation of pupation, we evaluated the impact of leafminer infestation on pupal development in the tomato mutant *def-1*. Importantly, while *def-1* plants are characterized by a reduced ability to produce jasmonic acid and to mount JA-regulated defense responses following mechanical damage and/or herbivory (Lightner *et al.*, 1993; Howe *et al.*, 1996), growth-related traits (e.g., plant height, number of leaves and dry mass) are unaffected (Li *et al.*, 2002; Thaler *et al.*, 2002; Escobar-Bravo *et al.*, 2017). Compared to non-infested wild type plants, leafminer pupae developed more rapidly in the soil of non-infested *def-1* plants (Fig 1bc, and Fig. S1, day 14, *def-1* non-infested vs wild type non-infested *p* ≤ 0.01, EMMeans pairwise comparisons). Strikingly, aboveground leafminer infestation of *def-1* plants reduced rather than increased adult emergence at 14 days after pupation (Fig. 1c, Tables S2, S3, and Fig. S1, *Genotype x Herbivory, χ* 23.03, df = 1, *p* < 0.01, ANOVA type II Wald chi-square tests; *def-1* non-infested vs *def-1* infested *p* ≤ 0.001, EMMeans pairwise comparisons). At the end of the experiment (day 16), no differences in adult emergence were observed between infested and non-infested *def-1* plants (Fig 1c, Tables S2, S3, and Fig. S1, *Genotype x Herbivory x Day, χ* 22.9, df = 1, *p* < 0.01, ANOVA type II Wald chi-square tests day 16, *def-1* non-infested vs infested *p* > 0.05, EMMeans pairwise comparisons). These results suggest that facilitation of pupal development is *def-1*- and, thus, likely jasmonate-dependent. They also demonstrate that the positive effect of aboveground herbivory on pupal development is plant-mediated rather than directly mediated by the feeding leafminer.

### Leafminer herbivory induces JA and ABA signaling pathways in the roots of wild type plants but not in *def-1*

To determine which plant systemic signals mediate leafminer facilitation of conspecific pupae in the soil, we measured the concentrations of defense-related hormones and the expression levels of hormonal marker genes in the leaves and roots of non-infested and infested wild type and *def-1* plants (Fig. 2).

**Figure 2.**
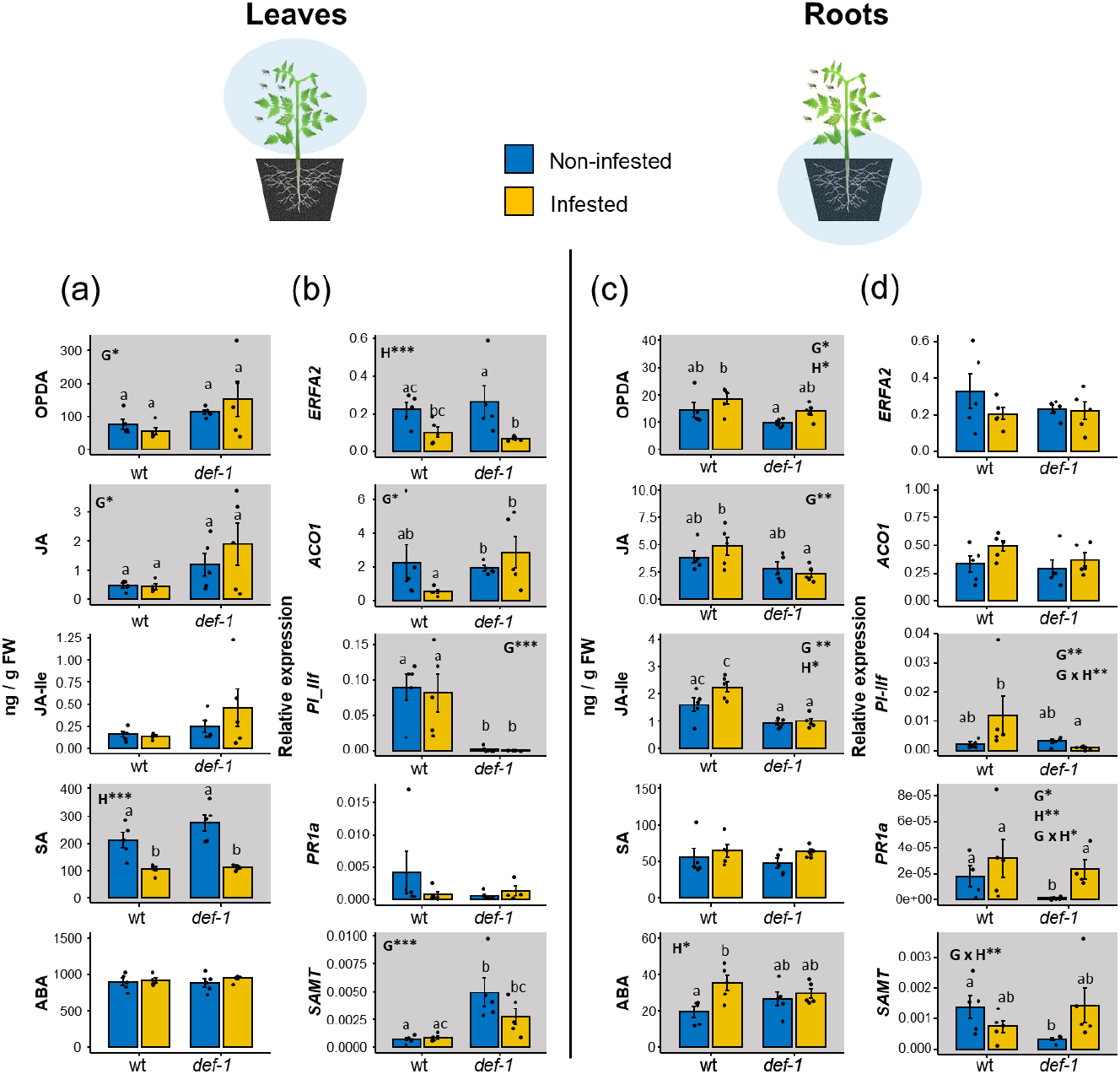
Aboveground leafminer infestation triggers different hormonal signaling in leaves and roots. Concentrations (mean ± SEM, *n* = 5) of 12-oxo-phytodienoic acid (OPDA), jasmonic acid (JA), jasmonic acid-isoleucine (JA-Ile), salicylic acid (SA), and abscisic acid (ABA) in leaves (**a**) and roots (**c**) of *L.trifolii-* infested and non-infested wild type (wt) and *def-1* plants at 24 days after infestation. Normalized transcript levels (mean ± SEM, *n* = 4-5) of marker genes for ethylene (*ERFA2* and *ACO1),* JA (*PI-IIf)* and SA (*PR-P1a*) signaling in leaves (**b**) and roots (**d**) of leafminer-infested and non-infested wt and *def-1* plants at 24 days after infestation. Transcript abundance were determined by qRT-PCR and normalized to *Ubiquitin* transcript levels. Shaded graphs indicate statistically significant effects of plant genotype (G), herbivory (H) and/or their interaction (G x H) determined by two-way ANOVAs (\**p* < 0.05, ** *p* < 0.01, *** *p* < 0.001). Different letters within plots indicate significant differences among groups tested by EMMeans post hoc test with multiple comparison adjustments (Tukey’s HSD) (*p* ≤ 0.05).

Constitutive levels of OPDA and JA were overall higher in the leaves of *def-1* plants compared to the wild type (ANOVAs, *genotype, p* < 0.05) (Fig. 2a,c), which is in line with previous reports (Howe et al., 1996; Li et al 2002). Levels of SA did not differ between both genotypes (Fig. 2a). Yet, we observed a higher expression of *SAMT* in *def-1* leaves (ANOVA, *genotype, p* < 0.001). This gene is responsible for the conversion of SA into methyl salicylate (MeSA) and therefore involved in the activation of systemic SA signaling. Upon leafminer infestation SA levels were significantly reduced in both wild-type and *def-1* leaves (ANOVA, *herbivory, p* < 0.001) (Fig. 2a). Leaf levels of OPDA, JA, JA-Ile and ABA were not affected by the leafminer infestation. Accordingly, the expression of the JA-responsive gene marker *PI-IIf* did not differ between non-infested and infested plants (Fig. 2b). Leafminer infestation did not affect the expression of the SA-responsive gene marker *PR1a* either, but it significantly reduced the expression of the ET-responsive marker *ERF2A* (ANOVA, *herbivory, p* < 0.001) (Fig. 2b). This suppression was not accompanied by a lower expression of the ET-biosynthetic related gene *ACO1*.

Levels of OPDA, JA and JA-Ile were higher in the roots of wild type plants than in *def-1* (ANOVAs, *genotype,p* < 0.05). Aboveground leafminer infestation overall increased OPDA, JA-Ile and ABA levels in wild type and *def-1* roots (ANOVAs, *herbivory, P* < 0.05) (Fig. 2c). When compared to their non-infested controls, however, differences in JA-Ile and ABA concentrations were slightly higher in the wild-type (JA-Ile, *p* = 0.06; ABA, *p* < 0.05; wt non-infested vs wt*-*infested, EMMeans pair-wise comparisons). *PI-IIf* expression levels were also higher in the roots of infested wild-type plants when compared to non-infested controls, and the opposite pattern was observed for *def-1* (ANOVA, *genotype x herbivory, p* < 0.01) (Fig. 2d). Besides jasmonates, SA levels were also higher in the roots of leafminer infested plants, albeit this was not statistically significant (ANOVA, *herbivory, p* > 0.05) (Fig. 2c). Relative expression of both the SA-responsive marker *PR1a* and SA-associated signaling *SAMT* genes, however, were significantly induced by leafminer attack in *def-1* only (ANOVA, *genotype x herbivory, p* < 0.05) (Fig. 2d). This suggests that there might be a stronger induction of SA signaling in the roots of the jasmonate deficient tomato mutant upon leafminer infestation. Together, these data reveal complex and non-uniform hormonal signaling induction in leaves versus roots and suggests that JA and ABA signaling are activated in roots of leafminer-infested plants in a genotype-dependent manner.

### Facilitation of pupal development is not dependent on the severity of leaf attack

The severity of plant damage upon herbivory can influence the magnitude of systemic plant responses, and, by consequence, plant-herbivore interactions (Robert *et al.*, 2012b). We thus tested whether leaf damage by *L. trifolii* correlates with pupal development in the soil of wild type and *def-1* plants. No significant difference in the number of leaf mines was found between wild type and *def-1* plants (Fig. 3a, t = 0.75, df = 17.86, *p* = 0.45, Student’s *t-*test). Similarly, no significant correlation between the number of leaf mines and the number of emerging adults at 14 days (Fig. 3b, *r* = - 0.08, *P* = 0.72, Pearson) nor at 16 days (*r* = - 0.092, *p* = 0.69, Pearson) was observed. Thus, facilitation of pupal development is unlikely to be driven by the severity of leaf attack, and the different effects observed in wild type and *def-1* plants are not explained by different leaf mining activities.

**Figure 3.**
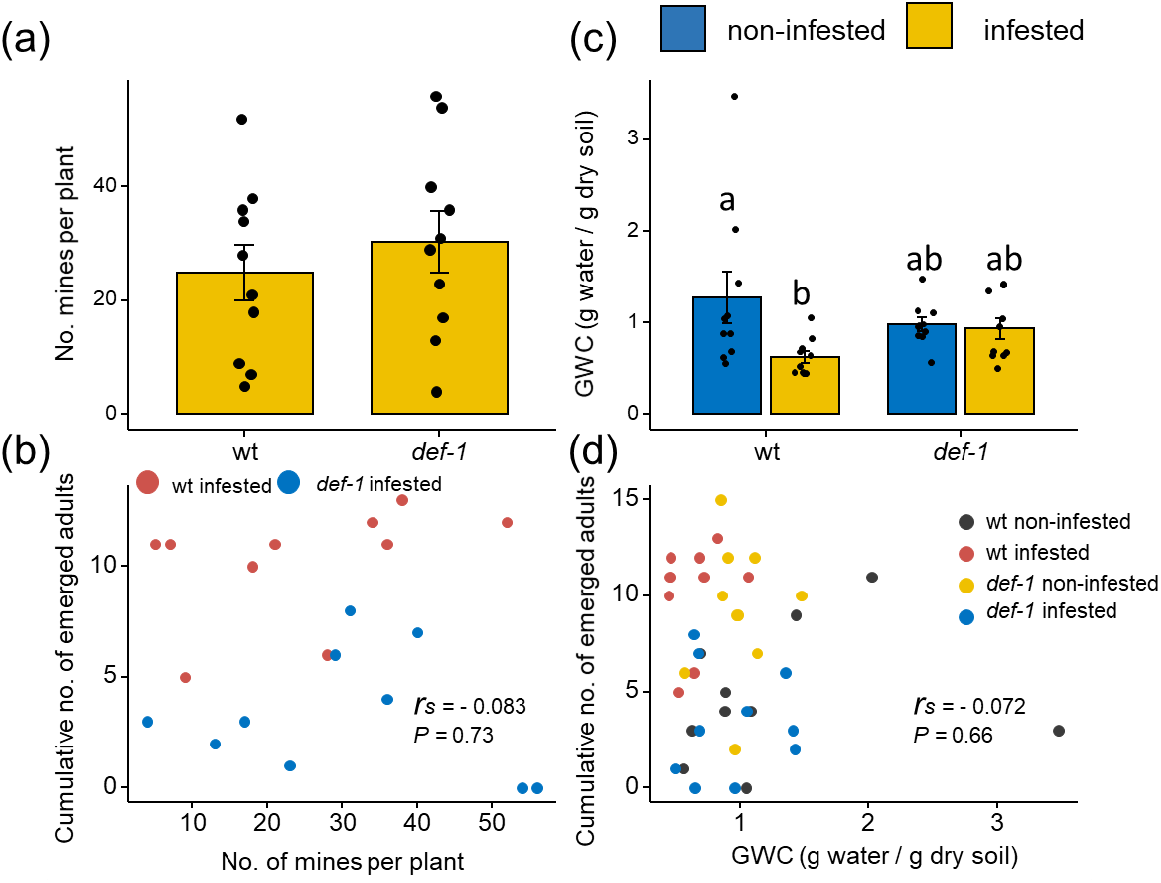
Facilitation of leafminer pupae in the soil is not correlated with leaf damage and soil humidity. (a) Number of mines per plant (mean ± SEM, *n* = 10) determined in wild type (wt) and *def-1* plants at 24 days after *L. trifolii* infestation. Plants were individually infested with seven *L. trifolii* adult female flies allowing them to feed and lay eggs into the leaves of the plant. No significant differences were found between treatments as tested by Student’s *t*-test at *p* ≤ 0.05. (b) Scatter plot depicting the relationship between the number of emerged *L. trifolii* adults at 14 days after pupation and the number of mines in *L. trifolii-*infested wt and *def-1* plants. The Pearson correlation coefficient (*r*) and *P* value are indicated. (c) Soil humidity (mean ± SEM, *n* = 10) estimated by using the gravimetric water content (GWC) method in non-infested and *L. trifolii-*infested wt and *def-1* plants at 24 days after infestation. Different letters indicate significant differences among groups tested by EMMeans post hoc test with multiple comparison adjustments (Tukey’s HSD) (*p* ≤ 0.05). (d) Scatter plot depicting the relationship between the number of emerged *L. trifolii* adults at 14 days after pupation and the soil humidity determined in non-infested and *L. trifolii-infested* wt and *def-1* plants at 24 days after infestation. The Spearman’s rank correlation coefficient (*r_s_*) and *p* value are indicated.

### Leafminer infestation changes soil humidity, but this does not explain facilitation of pupal development

Soil humidity is an important factor affecting pupal survival and adult emergence rate (Hou *et al.*, 2006; Chen & Shelton, 2007; Wen *et al.*, 2016). In most cases, extremely wet or dry soils can hinder adult emergence. We thus investigated whether *L. trifolii* leafminer attack changes soil humidity, and whether these changes are associated with the facilitation of pupal development. Leafminer attack significantly reduced soil humidity in wild type, but not in *def-1* plants (Fig. 3c, *Genotype, p* = 0.308, *Herbivory, p* < 0.05, *Genotype x Herbivory, p* = 0.0511, two-way ANOVA) (Fig. 3c). Differences in soil humidity were not correlated with the number of emerging adults at 14 (Fig. 3d, *r* = - 0.072, *p* = 0.66, Spearman) nor at 16 days (*r* = - 0.12, *p* = 0.44, Spearman). While these results cannot exclude the possibility that leafminer induced changes in soil humidity may have contributed to differences in pupal developmental times, they suggest that under our experimental conditions soil humidity does not play a major role in this process.

### Leafminer infestation modulates root volatile profiles in a *def-1* dependent manner

Leaf herbivory can change root volatile production (Robert *et al.*, 2012a; Huang *et al.*, 2017), which may influence pupal development. To gain first insights into the possible mechanism underlying the observed facilitation, we analyzed root volatile content in infested and non-infested wild type and *def-1* plants in a separate experiment. As leafminer infestation did not affect root biomass (Fig. S1, *genotype, p* = 0.14, *herbivory, p* = 0.42, *genotype x herbivory, p* = 0.14, two-way ANOVA), we expressed volatiles per g of root fresh weight. We detected 18 different volatile compounds in tomato roots. Redundancy analysis revealed that plant genotype, herbivory and their interaction explained 31.8 % of the total chemical variation (Fig. 4, F = 2.48, df = 3, *p* = 0.001, permutation *F-*test on the canonical *R^2^*). In the associated score plot, the first axis explained 51.1% of the total variation explained by the variables and separated wild type plants from *def-1* mutant plants. The second axis explained 38.7 % of the total variation and separated *L. trifolii* infested from non-infested wild type plants (Fig. 4a), with infested and-non infested *def-1* mutant plants clustering together in the middle. Permutation *F-*tests on individual variables confirmed this pattern and revealed significant differences in volatile profiles between plant genotypes and between infested and non-infested wild type plants, but not between infested and non-infested *def-1* plants (Fig. 4a).

**Figure 4.**
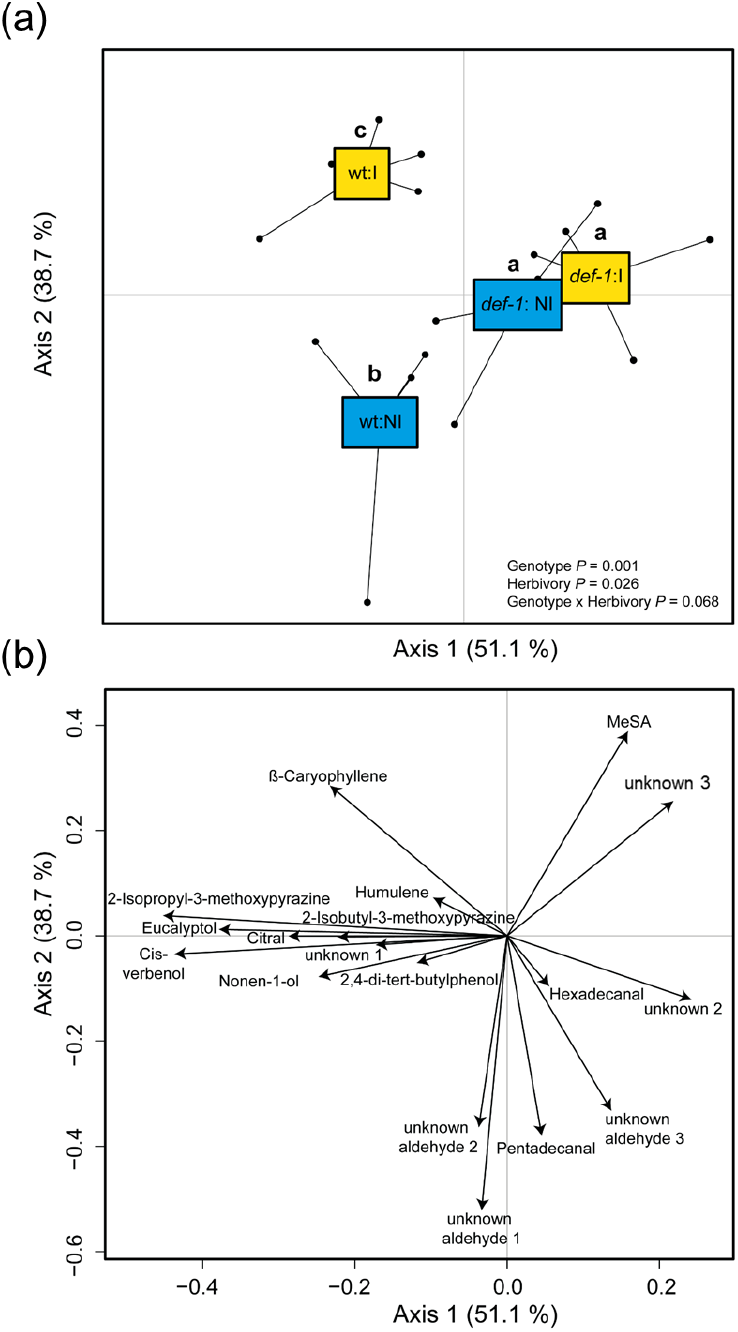
Leafminer herbivory modulates root volatile profiles in a *def-1* dependent manner. Redundancy analysis on tomato root volatile profiles analyzed by SPME-GC-MS in wild-type (wt) and *def-1* plants at 24 days after *L. trifolii* infestation (*n* = 5). (a) Score plot showing the first two principal axis with their explained variances in parentheses. Acronyms NI and I refers to non-infested and infested plants, respectively. The overall effects of plant genotype (G), herbivory (H), and their interaction (G x H) tested by permutation *F-*test are indicated. Different letters indicate significant differences among groups tested by permutation *F-*test pairwise comparisons corrected with the false discovery rate method at *p* ≤ 0.05. (b) Loading plot displaying the contribution of each volatile compound on the separation of the treatments. Vector’s length and direction denote the magnitude of their contribution and whether it is positive or negative, respectively.

Analysis of individual volatiles revealed that the levels of 1-8-cineole (eucalyptol), 2-isopropyl-3-methoxypyrazine, 2-isobutyl-3-methoxypyrazine, cis-verbenol and 3,7-dimethyl-2,6-octadienal (Citral) were significantly higher in wild type roots compared to *def-1* roots (ANOVAs, *genotype, p* < 0.05) (Fig. 4b and Fig 5). Leafminer infestation slightly increased the release of methyl salicylate in both genotypes (ANOVAs, *herbivory, p* < 0.05) (Fig. 5). Leafminer infestation suppressed the production of an unknown aldehyde #1 in the roots of wild type plants, but not *def-1* mutant plants (ANOVAs, *genotype x herbivory, p* < 0.05) (Fig. 5). A significant interaction between leaf infestation and genotype was also detected for (*E*)-β-caryophyllene (Fig. 5), which was induced by leaf infestation in the wild type but not in *def-1* plants (although pairwise comparisons were not significant for this compound). In summary, this experiment shows that leafminer infestation changes root volatile profiles in a *def-1* dependent manner, which is compatible with the hypothesis that systemic changes in root volatiles may be responsible for the observed changes in pupal developmental times.

**Figure 5.**
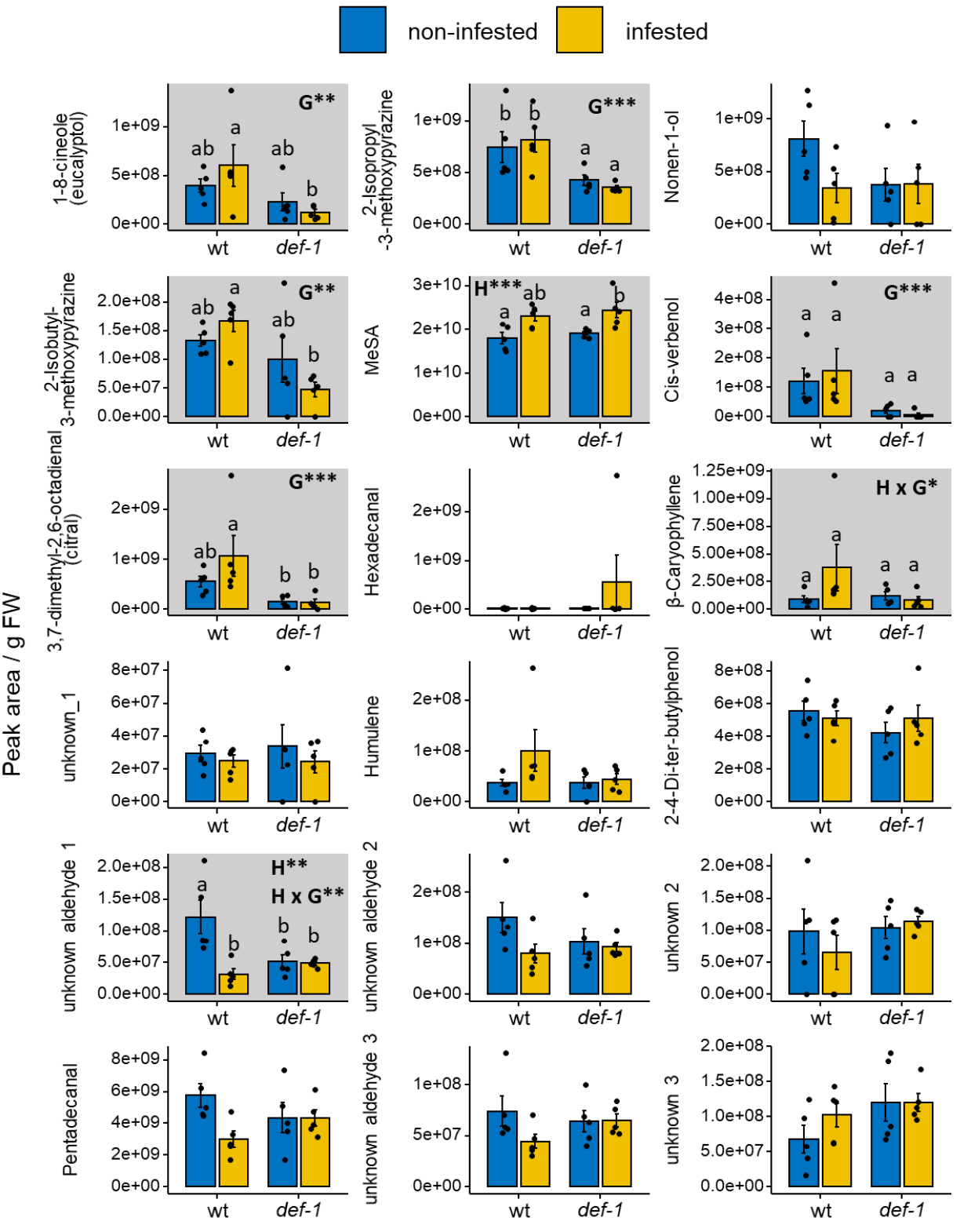
Leafminer herbivory affects the content of specific volatiles in tomato roots. Levels of tomato root volatiles (mean ± SEM, *n* = 5) detected in wild type (wt) and *def-1* plants at 24 days after *L. trifolii* infestation. Shaded graphs indicate statistically significant effects of plant genotype (G), herbivory (H) and/or their interaction (G x H) determined by two-way ANOVAs (\**p* < 0.05, ** *p* < 0.01, *** *p* < 0.001). Different letters indicate significant differences among groups tested by EMMeans post hoc test with multiple comparison adjustments (Tukey’s HSD) (*p* ≤ 0.05).

### Leafminer herbivory suppresses the expression of terpenoid biosynthetic genes in the leaves but not in the roots

To better understand how plant genotype and leafminer herbivory influence the production of root volatiles, we analyzed the expression levels of terpene-biosynthetic genes known to be involved in the production of the sesquiterpene β-caryophyllene and the monoterpene 1-8-cineole (Eucalyptol) (Fig. 6a) (Zhou and Pichersky, 2020) in the roots of non-infested and leafminer-infested plants. Levels of these compounds were significantly affected either by the plant genotype (Eucalyptol) or differently induced in the roots of infested wild type plants (β-caryophyllene) (Fig. 5). To determine the biological significance of these terpene biosynthetic genes in leafminer-tomato interactions, we also analyzed their expression levels in leaves. Constitutive levels of sesquiterpene- (*FPPS1* and *TPS12*) and some of the monoterpene- (*GGPPS3* and *CPT1*)-related biosynthetic genes were lower in the leaves of *def-1* plants compared to the wild type (ANOVAs, *genotype, p* < 0.05) (Fig. 6b). This might be explained by the reduced density of type-VI glandular trichomes, where these compounds are produced and stored, reported for this mutant (Pfeiffer et al., 2009, Escobar-Bravo et al., 2017). Leafminer infestation suppressed *GGPS1* expression (involved in the production of precursors for monoterpene biosynthesis) in the leaves of both genotypes (ANOVA, *herbivory, p* < 0.001). The same pattern was observed for *GGPS3, CPT1* and *SSU1,* albeit only statistically significant for *SSU1* and with a stronger suppression effect in *def-1* (ANOVA, *genotype* x *herbivory, p* < 0.01). In the roots, *SSU1* was less expressed in *def-1* plants than in the wild type (ANOVA, *genotype, p* < 0.05) (Fig. 6c), which mirrored the reduced levels of Eucalyptol detected in this genotype (Fig. 5). The expression pattern of *FPPS1* and *TPS12,* involved in β-caryophyllene biosynthesis, also mirrored the pattern of production of this compound in the roots of infested wild type and *def-1* plants, but this was not statistically significant (Fig. 5 and 6c).

**Figure 6.**
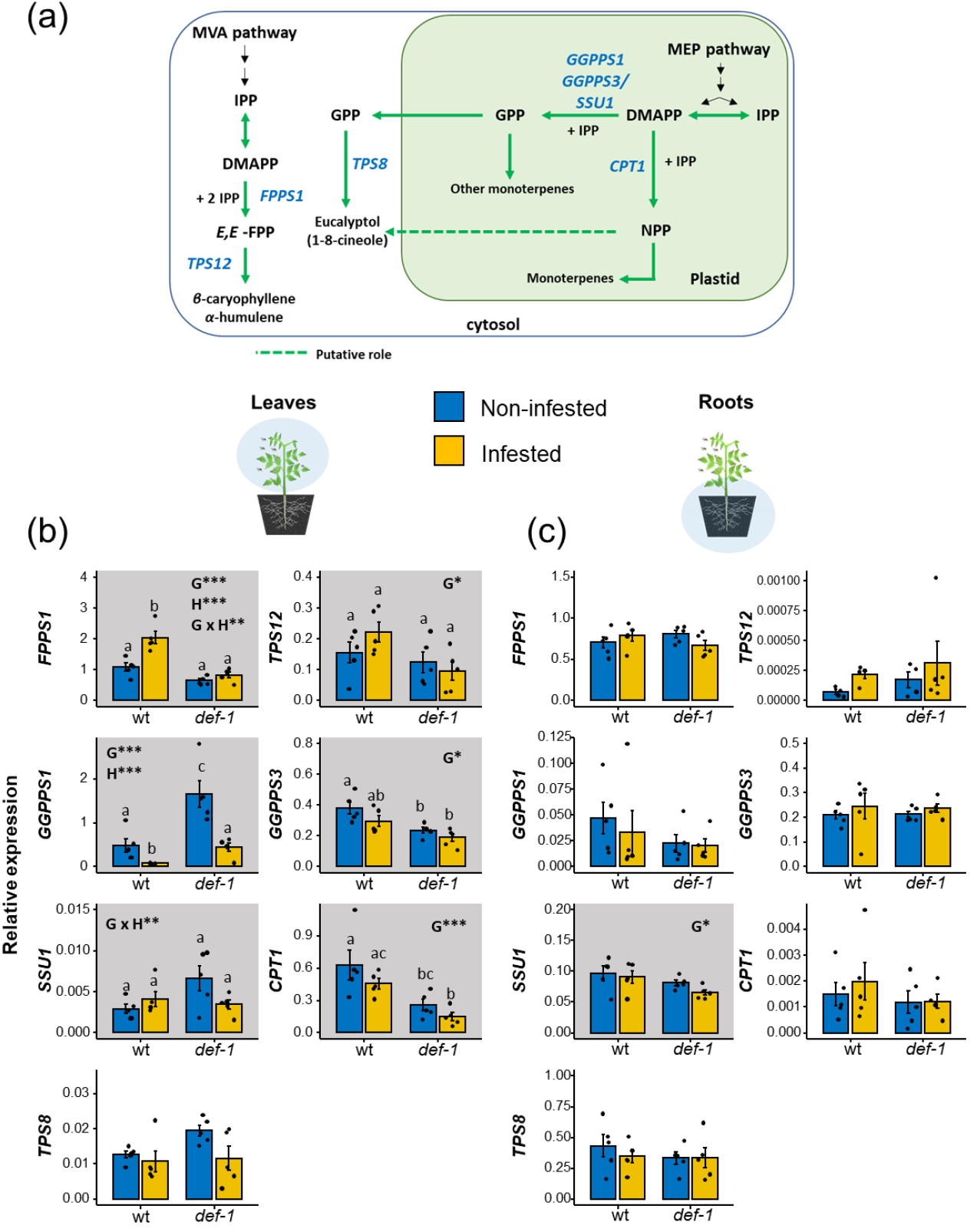
Leafminer herbivory effects on the expression of terpene biosynthetic genes in leaves and roots. **(a)** Simplified scheme displaying the biosynthesis pathway of β-caryophyllene and Eucalyptol (1,8-cineole) in tomato (adapted from Zhou and Pichersky, 2020). Terpene precursors are synthesized by the cytosolic mevalonic acid (MVA) pathway and the plastidic methylerythritol phosphate (MEP) pathway. Normalized transcript levels (mean ± SEM, *n* = 4-5) of terpene metabolism-related genes in **(b)** leaves and **(c)** roots of *L. trifolii*-infested and non-infested wild type (wt) and *def-1* plants at 24 days after infestation. Transcript abundance were determined by qRT-PCR and normalized to *Ubiquitin* transcript levels. Shaded graphs indicate statistically significant effects of plant genotype (G), herbivory (H) and/or their interaction (G x H) determined by two-way ANOVAs (\**p* < 0.05, ** *p* < 0.01, \*\*\**p* < 0.001). Different letters indicate significant differences among groups determined by EMMeans post hoc test with multiple comparison adjustments (Tukey’s HSD) (*p* ≤ 0.05). Abbreviations for precursors and genes: DMAPP, dimethylallyl diphosphate; IPP, isopentenyl diphosphate; GPP, geranyl diphosphate; NPP, neryl-diphosphate; *FPPS1, Farnesyl pyrophosphate synthase 1*; *TPS12, Terpene synthase 12*; *GGPPS1, geranylgeranyl diphosphate synthase 1*; *GGPPS3, geranylgeranyl diphosphate synthase 3*; *SSU1, small subunit of geranyl diphosphate synthase 1*; *CPT1, cis-prenyltransferase 1*; *TPS8, terpene synthase 8*.

## Discussion

Plants and herbivores interact dynamically with each other, with plants trying to mount appropriate defense responses and herbivores attempting to disrupt these responses and to modulate plant metabolism to their own benefit (Erb & Reymond, 2019). Here, we demonstrate that leafminer attack of aboveground plant tissues trigger systemic changes in the root signaling and metabolism that accelerates the development of the leafminer pupae in the soil adjacent to the plant. Below, we discuss this phenomenon from physiological and ecological points of view.

Over evolutionary time, herbivores have developed numerous strategies to feed and develop on well-defended plants, including the systemic manipulation of plant metabolism to improve their own performance and fitness (Sarmento *et al.*, 2011; Robert *et al.*, 2012a; Schimmel *et al.*, 2017; Xu *et al.*, 2019). Leaf damage by the red milkweed beetle *Tetraopes tetraophthalmus*, for instance, increases larval survival on the roots of the same plans (Erwin *et al.*, 2014). Similarly, aboveground feeding by adult *Bikasha collaris* beetles on *Triadica sebifera* enhances the survival of conspecific larvae on the roots while reducing the performance of heterospecific herbivores (Huang *et al.*, 2013; Huang *et al.*, 2014; Sun *et al.*, 2019). Whether insect herbivores can also modify plant metabolism to directly boost pupation -a critical step in their development- has not been investigated so far. The results reported here thus represent a new mechanism by which herbivores can facilitate the performance of their offspring. Faster pupal development likely increases *L. trifolii* performance by shortening its life cycle, thus potentially allowing for an increase in the number of generations per year. More importantly, faster pupal development may reduce the risk of predation by natural enemies in the soil such as entomopathogens (Liu, 2009). Both mechanisms may promote outbreaks of this pest species (Raymond *et al.*, 2002). Further work will reveal the importance of this form of plant-mediated facilitation for *L. trifolii* population dynamics and its interactions with other herbivore species and trophic levels. Many insect herbivores pupate on plants or in the soil close to plants and are thus exposed to plant chemicals. We therefore expect plant-mediated effects of herbivory on pupation success to be common in nature.

Herbivores may modulate the performance of other plant-associated herbivore species or life stages through different mechanisms, including direct (Kunert *et al.*, 2005) and plant-mediated effects (Soler *et al.*, 2012). Here, we present several lines of evidence in support of the second scenario. First, the modulation of leafminer pupal development by aboveground feeding conspecifics depends on the tomato genotype, with facilitation occurring in wild type plants and suppression in *def-1* mutants. *Def-1* is deficient in herbivory-induced JA accumulation (Li *et al.*, 2002) and defense induction, suggesting that the facilitation phenomenon observed in our study is jasmonate-dependent. Indeed, our data show that aboveground attack by the leafminer induced JA and ABA signaling in the roots of wild-type plants but not in *def-1.* This finding is in line with the central role of jasmonates and ABA in systemic signaling between leaves and roots (Baldwin & Zhang, 1997; Erb *et al.*, 2011; Machado *et al.*, 2013). We also found that induction of SA signaling by leafminer infestation in *def-1* roots coincided with delayed pupal development in the soil, suggesting a possible molecular mechanism for the reverse pattern in the tomato mutant. Second, leafminer attack triggers changes in soil-humidity and root volatile profiles in a *def-1* dependent manner, which is consistent with the notion that leaf attack modulates the physiology and metabolism of tomato roots in a way that facilitates pupal development in the soil. Finally, it should be kept in mind that leafminers, in contrast to other chewing herbivores, essentially live within the leaves. This means that they are unlikely to release high quantities of body odors and frass into the soil environment and influence their conspecific pupae (Hering, 1951). Taken together, these results strongly suggest that leafminers facilitate pupal development by inducing systemic changes in their host plant.

What is the eco-physiological mechanism by which tomato roots of leafminer infested plants facilitate pupal development in the soil? As the pupa in our experiments did not have direct root contact, changes in endogenous root chemistry can be excluded. On the other hand, soil-dwelling pupae are known to be highly sensitive to changes in abiotic conditions, such as soil humidity (Benoit *et al.*, 2010; Barnett & Johnson, 2013), and they may also be influenced by root exudate chemistry, including volatile and non-volatile root metabolites, which are known to modulate root herbivore behavior and performance (Rasmann *et al.*, 2005; Ali *et al.*, 2011; Robert *et al.*, 2012b; Robert *et al.*, 2012a; Huang *et al.*, 2017; Machado *et al.*, 2021). We found that leafminer infestation significantly reduced soil humidity in wild type plants, but not in *def-1* plants. Correlation analyses, however, did not show any association between soil humidity and pupal developmental speed. Furthermore, pupal development was slower in leafminer-infested than non-infested *def-1* mutants, even though soil humidity was not significantly affected by leafminer attack in this genotype. Thus, our data does not support a role of soil humidity in the systemic modulation of pupation time. Instead, we hypothesize that changes in exogenous root chemicals may be driving the observed patterns. In line with this, we observed that leafminer attack modulates root volatile production in a *def-1* dependent manner. Leafminer herbivory reduced the levels of an unknown aldehyde and slightly enhanced the accumulation of (*E*)-β-caryophyllene in wild type roots, but not in *def-1.* Although we did not find any volatiles that show a statistically significant inversion of responsiveness between the genotypes, tendencies for this phenomenon were found to occur for Eucalyptol and a isobutyl methoxypyrazine, two well-known plant aroma compounds (Lamy *et al.*, 2017; Murungi *et al.*, 2018), with Eucalyptol having antimicrobial properties (Kifer *et al.*, 2016). As belowground pupae are susceptible to fungal diseases, the differential emission of root volatiles may modulate the microbial load of the pupa, thereby indirectly altering their development. In addition, other belowground volatiles that were not detected in our analyses may have affected pupal development. For instance, belowground ethylene emissions can be modulated by aboveground herbivory (Robert *et al.*, 2012a), making this compound a target for future analyses. Our attempt to disentangle how leafminer attack affects the root chemistry at the molecular level showed that expression of terpene biosynthetic genes involved in the production of sesqui- and monoterpenes were overall suppressed in the leaves of wild type and *def-1* plants, but not affected in the roots. Yet, some genes involved in Eucalyptol biosynthesis mirrored the induced levels of this compound in the wild type upon leafminer infestation. Although wild type and *def-1* plants were continuously exposed to feeding leafminers during the experiments, a more detailed time-course analysis will shed light on the herbivory-mediated regulation of this metabolic pathway in the roots. Taken together, our work points to a role of systemic changes in root volatiles in mediating pupal development following leafminer attack. Further experiments with synthetic volatiles could unravel their specific impact on pupal development in the soil.

In conclusion, our study adds a novel facet to plant-herbivore interactions by demonstrating that leaf herbivory can facilitate pupal development in the soil. Plant volatiles likely play an important role in these interactions, as they are modulated by herbivore attack and readily reach non-herbivorous life stages of the herbivores, including pupa in the soil and on plants. This study represents a first step towards unravelling the molecular mechanisms and ecological significance of herbivory-induced modulation of metamorphosis.

## Acknowledgments

The work of BCJS was supported by Marie Sklodowska-Curie Action Individual Fellowship (European Union Horizon 2020, Grant Nr. 794,947).

## Author Contributions

REB conceived the study. REB and ME designed experiments. REB and BCJS performed experiments. REB and GG performed the hormone analyses. REB and BCJS performed gene expression analyses. REB analyzed the data. ME and PGLK supervised the study. REB, BCJS, ME and PGLK interpreted the data. REB wrote the manuscript and all authors contributed substantially to revisions.

## Data and materials availability

The data generated for this manuscript will be archived in Dryad and the data DOI will be included at the end of the article.

## Supplementary Material

**Table S1.**
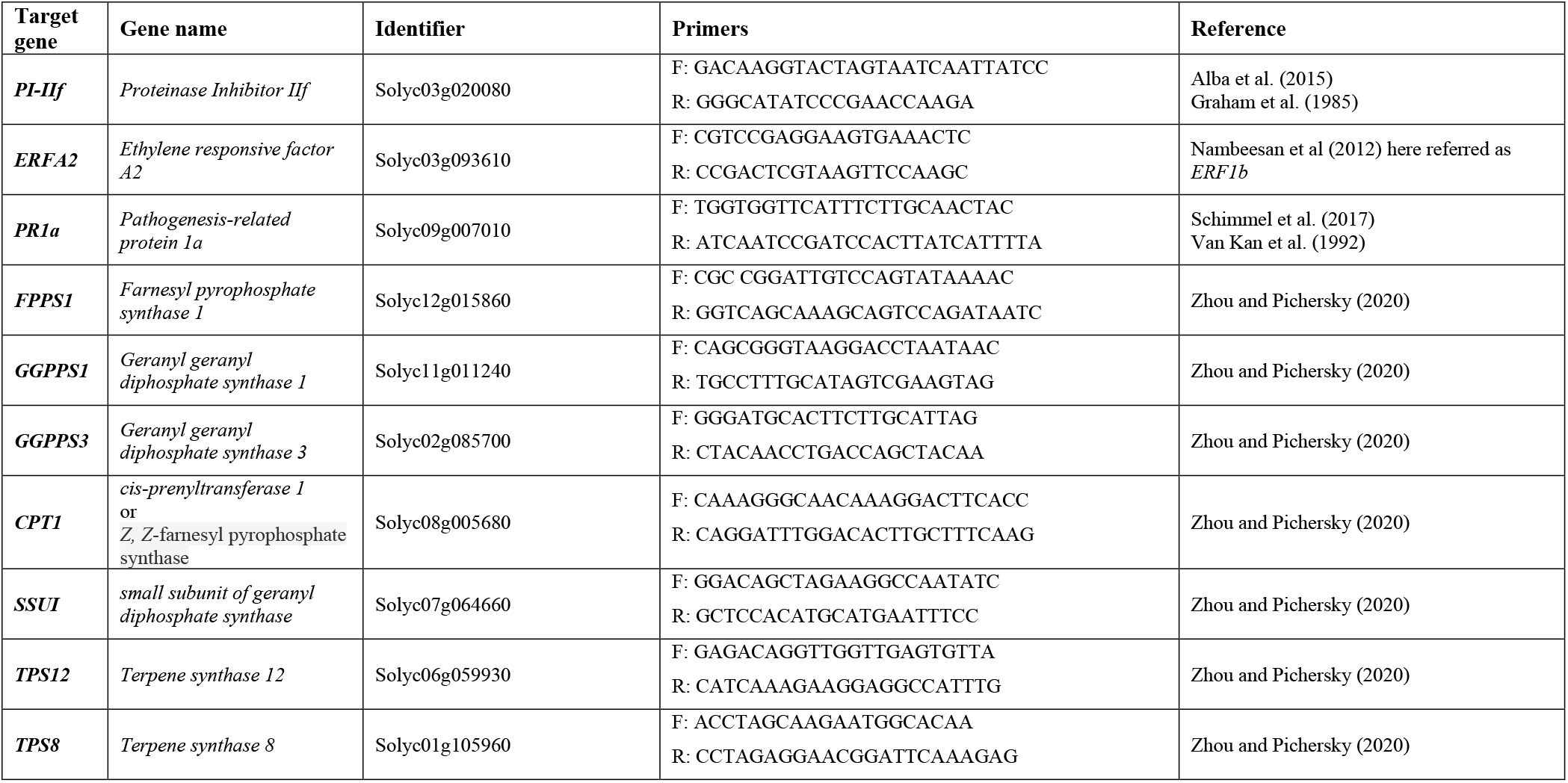
qRT-PCR primers specifications.

**Figure S1.**
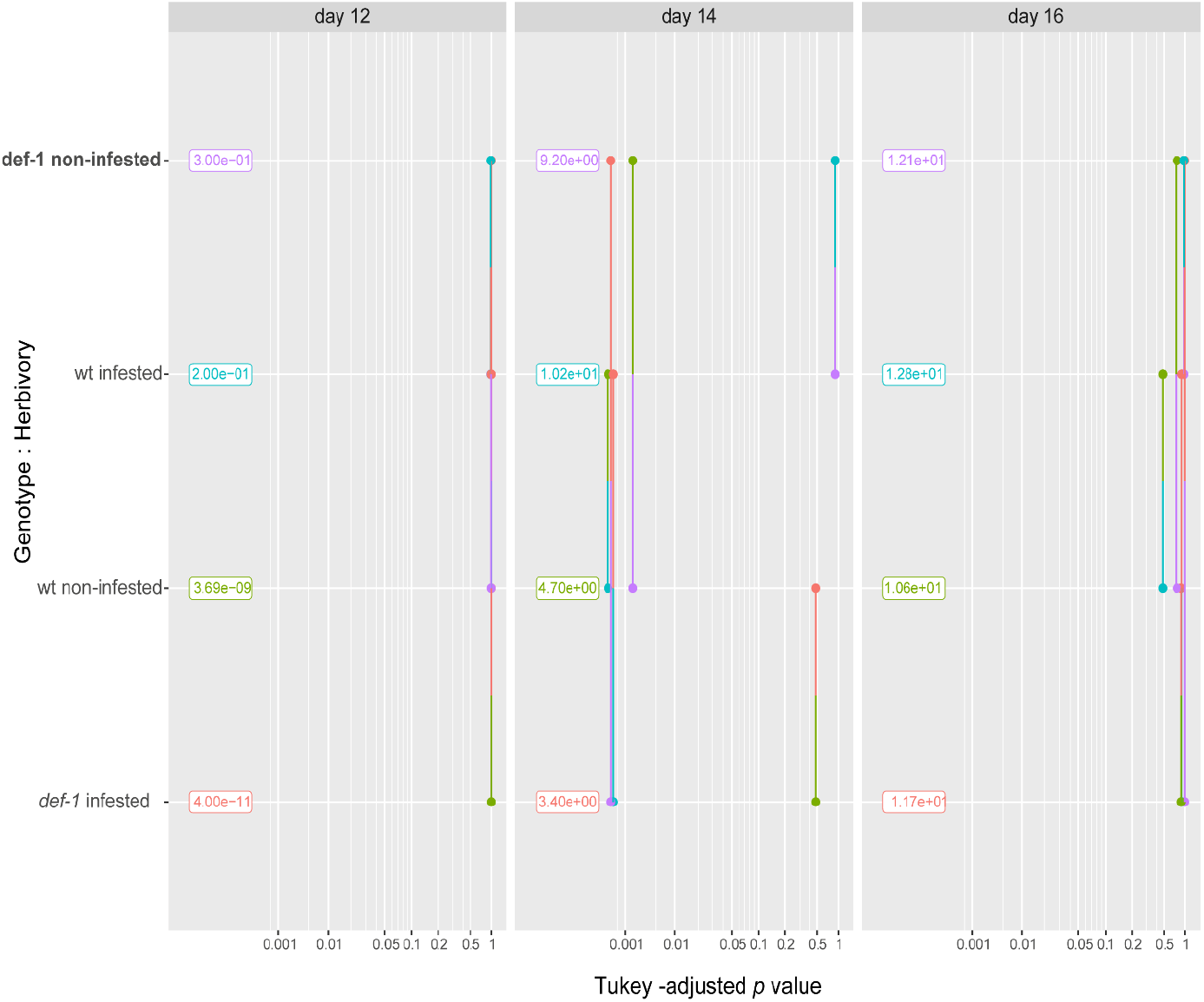
Plot of *p* values associated with pairwise comparisons of estimated marginal means. The effects of day of emergence, plant genotype, herbivory and their interactions, as well as the plant unit, on the cumulative number of emerged *L. trifolii* adults recorded in wild-type (wt) and *def-1* plants at 12, 14 and 16 days after pupation were modeled with a negative binomial model followed by pairwise comparisons of estimated marginal means.

**Table S2.** Analysis of deviance table (type II Wald Chi-square tests). The significance of the effects of day of emergence, plant genotype, herbivory and their interactions the cumulative number of emerged *L. trifolii* adults recorded in wild-type (wt) and *def-1* plants at 12, 14 and 16 days after pupation from the conditional model was tested via ANOVA-type analysis (type II Wald Chi-square tests).

| Factors | Chisq | Df | Pr (>Chisq) |
| --- | --- | --- | --- |
| Day** | 107.39 | 2 | $2.2 \cdot 10^{-16}$ |
| Genotype | 0.2060 | 1 | 0.6499 |
| Herbivory | 0.41 | 1 | 0.5207 |
| Day x Genotype | 1.61 | 2 | 0.44 |
| Day x Herbivory | 0.72 | 2 | 0.69 |
| Genotype x Herbivory** | 23.03 | 1 | $1.59 \cdot 10^{-06}$ |
| Day x Genotype x Herbivory** | 22.90 | 1 | $1.06 \cdot 10^{-05}$ |

\*\* $p < 0.001$

**Table S3.** Estimated marginal means, standard errors and confidence limits.

| Day | Genotype | Herbivory | Response | SE | Df | Lower CL | Upper CL |
| --- | --- | --- | --- | --- | --- | --- | --- |
| 12 | *def-1* | infested | $1.20 \cdot 10^{-05}$ | 2.00e-06 | 107 | 0.0 | $1.02 \cdot 10^{50}$ |
| 14 | *def-1* | Infested | 3.4 | 5.83e-01 | 107 | 2.42 | 5 |
| 16 | *def-1* | Infested | 11.7 | 1.08e+00 | 107 | 9.73 | 14 |
| 12 | wt | Infested | 0.2 | 1.41e-01 | 107 | 0.04 | 1 |
| 14 | wt | Infested | 10.2 | 1.01e+00 | 107 | 8.38 | 12 |
| 16 | wt | Infested | 12.8 | 1.13e+00 | 107 | 10.75 | 15 |
| 12 | *def-1* | Non-infested | 0.3 | 1.73e-01 | 107 | 0.09 | 1 |
| 14 | *def-1* | Non-infested | 9.2 | 9.59e-01 | 107 | 7.47 | 11 |
| 16 | *def-1* | Non-infested | 12.1 | 1.10e+00 | 107 | 10.11 | 14 |
| 12 | wt | Non-infested | 0.0 | 1.92e-05 | 107 | 0.00 | $7.58 \cdot 10^{13}$ |
| 14 | wt | Non-infested | 4.7 | 6.86e-01 | 107 | 3.51 | 6 |
| 16 | wt | Non-infested | 10.6 | 1.03e+00 | 107 | 8.74 | 13 |

Confidence level used: 0.95
Intervals are back transformed from the log scale

**Figure S2.**
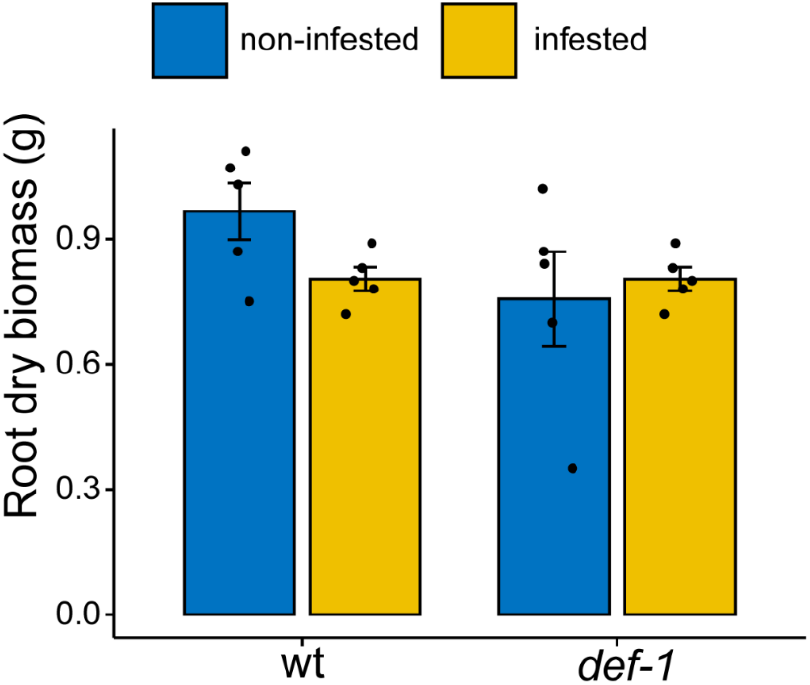
Effect of aboveground *L. trifolii* herbivory root dry biomass. Average root dry biomass (mean ± SEM, *n* = 5) determined in non-infested and infested wild-type (wt) and *def-1* plants at 24 d after *L. trifollii* infestation.

## Notes

### Competing Interest Statement

The authors have declared no competing interest.

